# Mass spectrometry-based multi-omics identifies metabolic signatures of sarcopenia in rhesus monkey skeletal muscle

**DOI:** 10.1101/2023.07.31.551315

**Authors:** Melissa R. Pergande, Katie J. Osterbauer, Kevin M. Buck, David S. Roberts, Nina N. Wood, Priya Balasubramanian, Morgan W. Mann, Kalina J. Rossler, Gary M. Diffee, Ricki J. Colman, Rozalyn M. Anderson, Ying Ge

## Abstract

Sarcopenia is a progressive disorder characterized by age-related loss of skeletal muscle mass and function. Although significant progress has been made over the years to identify the molecular determinants of sarcopenia, the precise mechanisms underlying the age-related loss of contractile function remains unclear. Advances in ‘omics’ technologies, including mass spectrometry-based proteomic and metabolomic analyses, offer great opportunities to better understand sarcopenia. Herein, we performed mass spectrometry-based analyses of the vastus lateralis from young, middle-aged, and older rhesus monkeys to identify molecular signatures of sarcopenia. In our proteomic analysis, we identified numerous proteins that change with age, including those involved in adenosine triphosphate and adenosine monophosphate metabolism as well as fatty acid beta oxidation. In our untargeted metabolomic analysis, we identified multiple metabolites that changed with age largely related to energy metabolism including fatty acid beta oxidation. Pathway analysis of age-responsive proteins and metabolites revealed changes in muscle structure and contraction as well as lipid, carbohydrate, and purine metabolism. Together, this study discovers new metabolic signatures and offer new insights into the molecular mechanism underlying sarcopenia for the evaluation and monitoring of therapeutic treatment of sarcopenia.

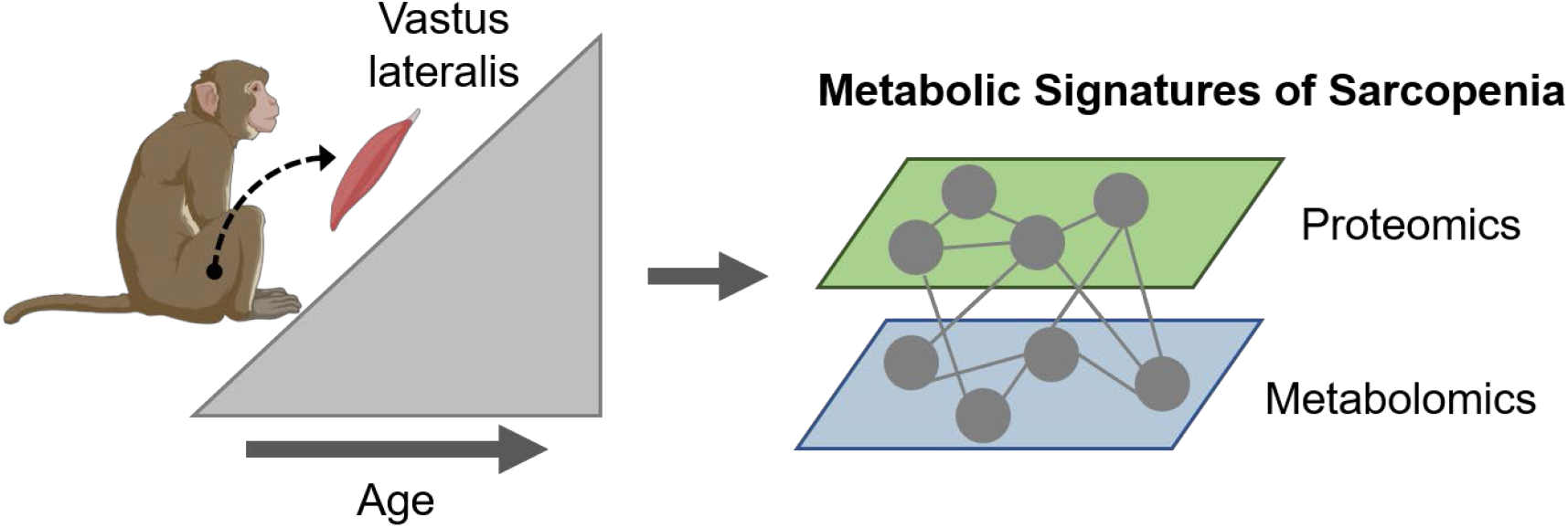

Graphical Abstract

## Introduction

Sarcopenia is the loss of skeletal muscle mass and function that occurs during aging which leads to muscle weakness, falls, fractures, and hospital admissions in the elderly and represents an increasingly serious public health problem.^1–5^ A major hallmark of sarcopenia is the progressive decline in skeletal muscle contractile function. Unfortunately, the molecular mechanisms underlying this loss of function remain poorly understood.^1, 4–6^ Previous reports have suggested that the lower contractile capacity observed with age is due, in part, to alterations in the structure and function of myofilaments,^7^ major protein components of the sarcomere which are responsible for force production during muscle contraction.^8–10^ In other studies, it has been reported that loss of contractile function is directly associated with an age-related decline in oxidative capacity, suggesting mitochondrial dysregulation as a major factor underlying sarcopenia.^11–16^ Despite these findings, there remains a need to identify the precise molecular determinants of contractile dysfunction to attenuate or prevent loss of muscle mass and function in aging individuals.

Non-human primates (NHPs), in particular rhesus monkeys, have shown to be a valuable and highly translational research model for studying sarcopenia.^17–19^ Understanding the molecular mechanisms that lead to age-related loss of muscle mass and contractile dysfunction in primates is further complicated by the observation that no two individuals age in the same manner or at the same rate. Many biological models including cells, yeast, worms, flies, zebrafish, and rodents have provided a basic understanding of age-related molecular changes; however, these models lack the complexity of human aging.^20–22^ Rhesus monkeys share ∼93% genome homology with humans,^23, 24^ likely contributing to their marked similarity in age-related diseases and pathophysiology.^25–27^ Thus, monkeys provide the best opportunity to study the underlying age-related molecular perturbations which contribute to human sarcopenia.

Mass spectrometry (MS)-based analyses allow the opportunity for an in-depth understanding of diseases at a molecular level, providing the foundations for systems biology approaches. MS-based ‘omic’ technologies specifically provide an unbiased means to profile the molecular landscape in complex biological systems.^28–30^ Unlike genes, proteins and metabolites are closer to the endpoint of the ‘omics cascade, and provide a different perspective on disease phenotype. When proteomic and metabolomic data are acquired from the same samples, the interplay between protein and metabolite levels can be directly evaluated. It is important to note that protein-metabolite dynamics are very complex and this approach does not consider non-linear kinetic mechanisms, substrate affinity, enzyme activity, metabolite-metabolite negative or positive feedback, or post-translational modifications.^31, 32^ Therefore, a multi-omic approach can provide the opportunity to identify relationships among groups of molecules and how they work in concert in the setting of disease pathophysiology.^33–38^

In this study, we conducted a mass spectrometry-based multi-omics analysis of the vastus lateralis from young adult, middle-aged, and older age monkeys after biometric and histological characterization of muscle structure and quality. Next, we performed an unbiased analysis, defining proteomic and metabolomic profiles and integrated those data to identify concerted sarcopenia-associated perturbations. The results of this study provide insight regarding age-related metabolic alterations and identifies potential targets for therapeutic treatment.

## Materials and Methods

### Chemicals and reagents

All reagents were purchased from Millipore Sigma (St Louis, MO, USA) and Fisher Scientific (Fair Lawn, NJ, USA) unless noted otherwise.

### Animals, biometrics, and biopsies

This study involved male young adult (median age 7.2y), middle-age (median age 15.1y), and older (median age 28.1y) rhesus macaques (*Macaca mulatta*) that were born and housed at the Wisconsin National Primate Research Center, and maintained according to protocols approved by the Institutional Animal Care and Use Committee of the University of Wisconsin-Madison. Body weight and height (crown-to-rump) taken at time of biopsy collection were used to calculate BMI. Electrical properties of the muscle of the upper leg were determined using segmental bioelectrical impedance spectroscopy (S-BIS). Bioelectrical impedance was measured using a logarithmic distribution of 256 frequencies, ranging from 4 to 1,000 kHz (SFB7, ImpediMed, Pinkenba, QLD, Australia), using disposable tab-type monitoring electrodes (2 cm x 2 cm, Medtronic, Minneapolis, MN, USA). The R0 and R∞ for the whole-body and leg were determined by extrapolation after fitting the spectrum of bioimpedance data to the Cole-Cole model and the RI was calculated using 1/[(1/R∞) − (1/R0)]. Segment length (L; cm) was measured from the most lateral aspect of the lateral greater trochanter of the femur to lateral tibial malleolus. Intracellular impedance index was calculated as L2/RI, and extracellular resistance index was calculated as L2/R0. L2/RI reflects segmental ICW, and L2/R0 reflects segmental ECW in the leg. The resistance ratio was calculated as R0/RI, which is the index of the ratio of ICW/ECW. The membrane capacitance (Cm), characteristics frequency (fc), and phase angle (φ) were also obtained from the Cole-Cole model.^39^ Vastus lateralis skeletal muscle was collected, bisected, and either flash frozen in liquid nitrogen or embedded in Optimal Cutting Temperature Medium (OCT, Sakura Inc., Torrance, CA) and frozen in liquid nitrogen. Samples were stored at -80 °C until use.

### Histology

For histochemical analysis all specimens were processed simultaneously for each stain as previously described.^16, 40^ Slide sections were stained with hematoxylin and eosin for non-contractile (i.e., non-myofibillar) content quantification. Fibrosis was detecting using picrosirius staining (Sigma Cat#365548) according to manufacturer’s instructions. For fiber typing, fiber type specific expression of myosin chain isoforms was detected using specific antibodies (M8421, skeletal, slow myosin, monoclonal clone NOQ7.5.4D, Sigma-Aldrich, St. Louis, MO; M1570, skeletal, fast myosin, monoclonal clone MY-32, Sigma-Aldrich).

### Proteomic Analysis

Approximately 5 mg of skeletal muscle tissue from each monkey was suspended in 500 μL of tissue lysis buffer (4% SDS, 100 mM Tris HCl pH 7.6, 100 mM DT containing 1x Halt Protease inhibitor cocktail), homogenized using a Teflon pestle, and heated for 5 min at 96 °C.^41^ Each of the homogenates were subjected to three rounds of probe sonication (20% amplitude for 10 s with alternating bursts every 1 second). The resulting tissue lysates were centrifuged for 10 min at 16,100 *g* (Sorvall Legend Micro 21R, Thermo Fisher Scientific, Am Kalkbarg, Germany). Next, the supernatant was removed and protein concentration determined via a bicinchoninic acid assay. Thirty micrograms of protein from each sample was taken for further processing. Briefly, samples were reduced with 10mM of dithiothreitol for 15 min at 55°C and alkylated with 30mM chloroacetic acid for 20 min at room temperature in the dark. Enzymatic digestion and detergent removal were performed using the S-Trap device and Trypsin Gold (Promega, Madison, Wisconsin, USA) following the manufactures recommended protocol. The dried peptides were reconstituted in 50 µL of 0.1% formic acid. Protein concentration was normalized to 0.2 µg/µL using a NanoDrop One Microvolume UV-Vis Spectrophotometer (Thermo Fisher Scientific, Waltham, MA, USA) to determine peptide concentration through A205 response.

Analysis of the protein digests were performed using a nanoElute liquid chromatography (LC) system (Bruker, Bremen, Germany) coupled to the timsTOF Pro (Bruker, Bremen, Germany) mass spectrometer (MS) where mobile phase A was 0.2% formic acid and mobile phase B was acetonitrile containing 0.2% formic acid. For each sample, 250 ng was loaded on a capillary C18 column (25cm length, 75µm inner diameter, 1.6 µm particle size, 120 Å pore size; IonOpticks, Fitzroy, VIC, AUS). Peptides were separated using a 90-min gradient at a flow rate of 400 nL/min (0-60’ 2-17%B, 60-90’ 17-25%B, 90-100’ 25-37%B, 100-110’ 37-85%B, 110-120’ 85%B) at 55°C. MS spectra were collected in data independent parallel accumulation serial fragmentation (diaPASEF) mode, using 32 windows ranging from *m/z* 400 to 1200 and 1/K0 0.6 to 1.42.

Raw LC-MS global proteomics data was processed using DIA-Neural Network (DIA-NN)^42^ using the default parameters: 1% FDR, library-free search enabled, *m/z* 300-1800 precursor range, *m/z* 200-1800 fragment ion range, 2-4 precursor charge range, 7-30 peptide length, 2 maximum missed cleavages, robust LC (High Precision) quantification strategy, and double-pass neural network classifier mode. A library was generated by using the reviewed UniProtKB/Swiss- Prot *Homosapien* (20,423 sequences) database with Deep learning-based spectra, RTs and IMs prediction selected. Trypsin/P was set as the protease with two missed cleavages and carbamidomethyl (C) was set as fixed modifications cleavages and searches were performed with a fragment mass error tolerance of ≤10 ppm with a minimum of one unique peptide. Protein-level quantification was performed using the DAPAR^43^ R package for proteins identified in 3 of 4 biological replicates in at least one sample group. Intensity values were median normalized and missing values were imputed via ssla for partially observed values within a sample group or set to the 2.5% quantile of observed intensities for observations that were missing entirely within a condition. The DEP R package was used to perform a Limma test between all specified contrasts.^44^ The IHW R package was used to adjust all p-values, using the number of quantified peptides per protein as a covariate.^45^ An adjusted p-value (p ≤ 0.05) was used to determine significant changes to protein expression. Protein intensities for selected proteins were plotted for visual comparison using GraphPad Prism (v9.4.1). Any missing values were imputed using the average intensity within a sample group and significance was displayed using an ordinary one-way ANOVA with multiple comparisons (Old vs Young, Old vs Middle, and Middle vs Young).

### Metabolomic Analysis

Ten milligrams (+/-0.1 mg) of rhesus monkey skeletal muscle tissue were homogenized in 500 μL of ice-cold (-20 °C) methanol using a Teflon pestle, vortexed for 10 s, and incubated for 15 min at 4°C to extract metabolites. The resulting homogenate was centrifuged for 5 min at 14,000*g*, supernatant removed, and extracts dried *in vacuo*. Metabolites were reconstituted in 100µL of 75% methanol and analyzed in both positive and negative ion modes using LC-MS by data dependent acquisition where mobile phase A was 6:4 acetonitrile: water containing 10 mM ammonium formate and 0.1% formic acid and mobile phase B was 9:1 2-propanol: acetonitrile containing 10mM ammonium formate and 0.1% formic acid. For negative ion mode, ammonium acetate was used as mobile phase additive instead of ammonium formate and formic acid. Metabolites were separated on a nanoEase M/Z HSS T3, 100mm length x 300μm id, 1.8μm particle size, 100Å pore size column (Waters, Milford, MA, USA) by means of a 20-min gradient (0-5’ 60-82% B, 5-15’ 82-100% B, 15-17’ 100% B, 17.0-17.1 60% B, 17.1-25’ 60%B) using a nanoACQUITY UPLC system (Waters, Milford, MA, USA). Mass detection was performed using an Impact II quadrupole-time-of-flight mass spectrometer (Bruker, Bremen, Germany) where mass spectra were collected at 5Hz over a *m/z* 50-1200 range.

Raw LC-MS/MS metabolomics data was processed using MetaboScape (v2022b). Bucket lists for data collected in positive and negative ionization modes were generated using the T-Rex 3D algorithm and combined to create one merged bucket list using a mass error and retention time tolerance of ≤5.0ppm and 20 seconds, respectively. Metabolite annotations were performed by searching tandem MS data against publicly available databases from MassBank including NoNA-export-LipidBlast, NoNA-export-Experimental_Spectra, NoNA-export-HMDB, NoNA- export-MassBank, NoNA-export-LC–MS-MS_Spectra databases with an error tolerance ≤10.0 ppm. Metabolomics statistical analysis was performed comparing peak areas in the MetaboScape software using a t-test between groups where an adjusted p-value (p ≤ 0.05) was used to determine significant changes to metabolite abundance. Selected metabolites were plotted for visual comparison using GraphPad Prism (v9.4.1, San Diego, CA, USA) where significance was displayed using an ordinary one-way ANOVA with multiple comparisons (Old vs Young, Old vs Middle, and Middle vs Young).

### Gene Ontology Enrichment and Pathway Analysis

Gene Ontology (GO) enrichment analysis was performed on altered proteins between age groups to identify associated biological processes. Protein names were copied into the GO search tool where *Homo sapiens was used* for the enrichment analysis. Only results with a *p* ≤ 0.05, calculated by the Mann–Whitney *U* test, were used. Pathway analysis of dysregulated proteins and metabolites was performed. Here, significantly altered up and downregulated proteins and metabolites were uploaded to MetaboAnalyst 5.0 for integrative pathway analysis where ranking of enriched pathways was determined by using a Fisher’s exact test for Enrichment analysis, Degree Centrality for Topology measure, and Combine p-values (pathway level) for the Integration method.^46^ The output from the pathway analysis was plotted GraphPad Prism (v9.4.1) where selected pathways were annotated.

## Results and Discussion

In this study, we conducted a MS-based multi-omic analyses of skeletal muscle from young adult, middle-aged, and older age monkeys after biometric and histological characterization of muscle structure and quality to identify molecular signatures of sarcopenia in aging rhesus monkeys (**Figure 1**). Here, we chose vastus lateralis as it is a mixed fiber type muscle (comprised of Type I and Type II myosin expressing fibers) and in primates it is easily accessible, making it an excellent choice for cellular and molecular studies of sarcopenia.^19^ Moreover, the impact of age in humans and in monkeys is fiber type specific, and fibers expressing Type II myosin are more vulnerable to age-related loss and function.^16, 47, 48^

**Figure 1.**
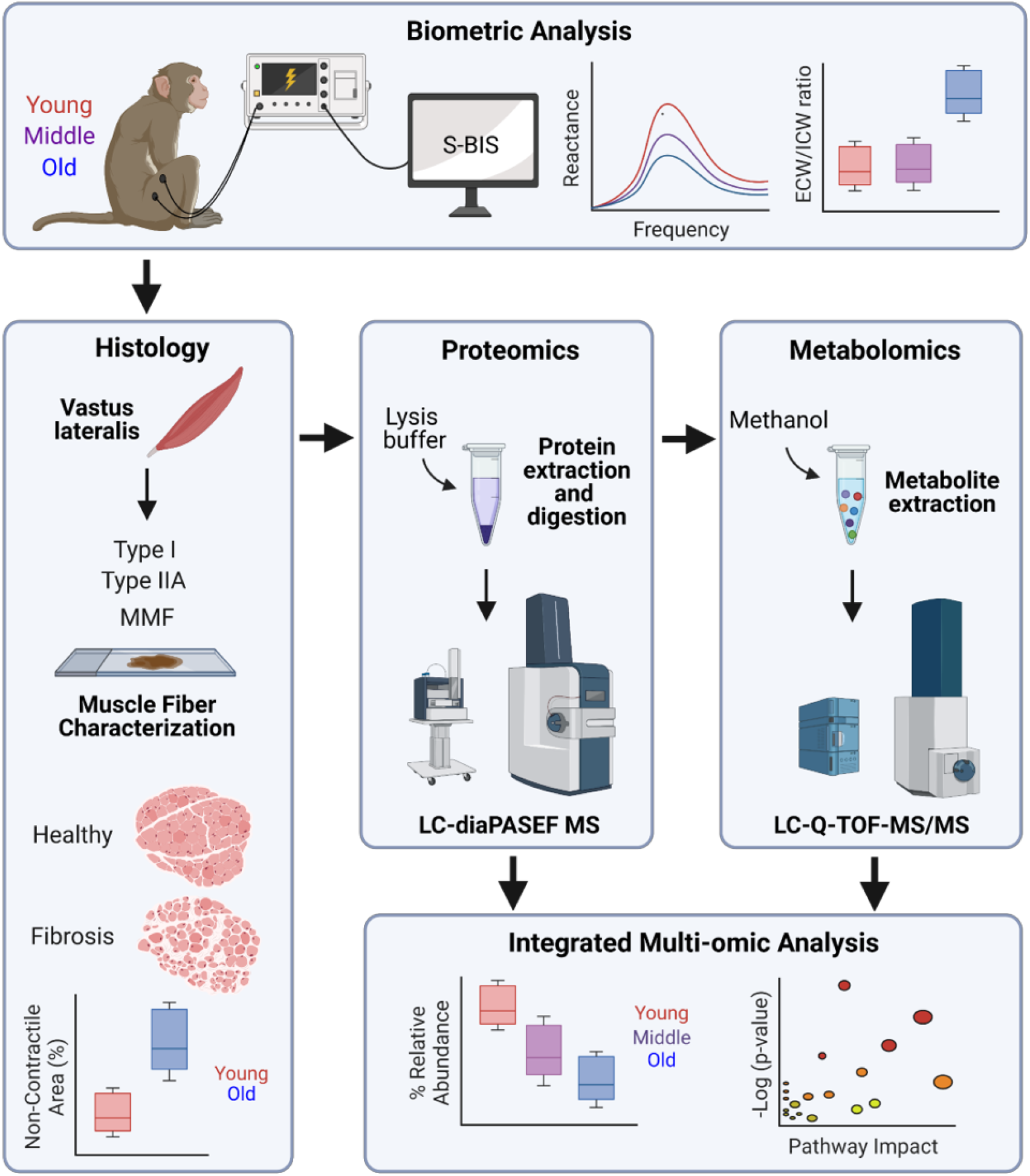
Overview of the integrated functional and multi-omic analysis of aging rhesus monkey skeletal muscle. Biometric analysis including segmental bioelectrical impedance spectroscopy (S-BIS) was performed on the vastus lateralis from young (median age 7.2y), middle-age (median age 15.1y) and old (median age 28.1y) rhesus monkeys (n=4 per group). Histological measurements including fiber type and non-contractile area characterization was performed. After protein extraction and enzymatic digestion, proteome changes were measured using LC-data independent parallel accumulation serial fragmentation mass spectrometry (LC-diaPASEF MS). An untargeted metabolomics study was performed to measure altered metabolites between groups by LC-Q-TOF tandem MS (MS/MS) analysis after extraction of metabolites. An integrated multi-omics analysis was performed of significantly altered proteins and metabolites to identify concerted biological changes. Figure created with *BioRender.com*.

### Biometric and Histology Analyses

This study involved male adult young (median age 7.2y), middle-age (median age 15.1y), and old (median age 28.1y) rhesus monkeys-age groups that represent three phases across the lifespan: young adult, healthy stage when monkeys are full stature, middle-age adult the time of sarcopenia onset but before there are signs of pathology, and advanced age when sarcopenia is overt (**Figure 2A**).^19^ As a reference, median lifespan in captive rhesus monkeys is ∼26 years, 10% survival is ∼35 years, maximum lifespan is ∼40 years.^49^ Bodyweight and body mass index (BMI: kg/m^2^) were both age-sensitive with non-linear changes over life course, although the differences were not statistically different (**Figure 2B**). Age-related changes in muscle composition were captured using in vivo S-BIS. The electrical properties of muscle correlate with functional capacity in humans, including its decline with age, even independent of loss of muscle mass.^39^ Electrical resistance measures were used to calculate extracellular/intracellular water (ECW/ICW) ratio that was higher in animals of advanced age (**Figure 2C**). The phase angle, (derived from Cole-Cole plots) was significantly lower in older animals and reactance as a function of frequency applied showed age-related changes (**Figure 2C**). These differences in electrical properties are consistent with changes in the underlying composition of muscle, and indicate that the advanced age monkeys were sarcopenic. Histological analysis reveals changes in fiber type distribution along with increased heterogeneity in fiber cross sectional area (**Figure 2D**), all of which are established features of skeletal muscle aging. Percent non-myofiber area increased significantly with age along with fibrosis, with numerically higher intensity of picrosirius stain in biopsy sections from older animals (**Figure 2E**). These data show that overt pathology and tissue level biomarkers of skeletal muscle aging are absent in the middle-age animals, and confirm overt sarcopenia in the advanced-age animals.

**Figure 2.**
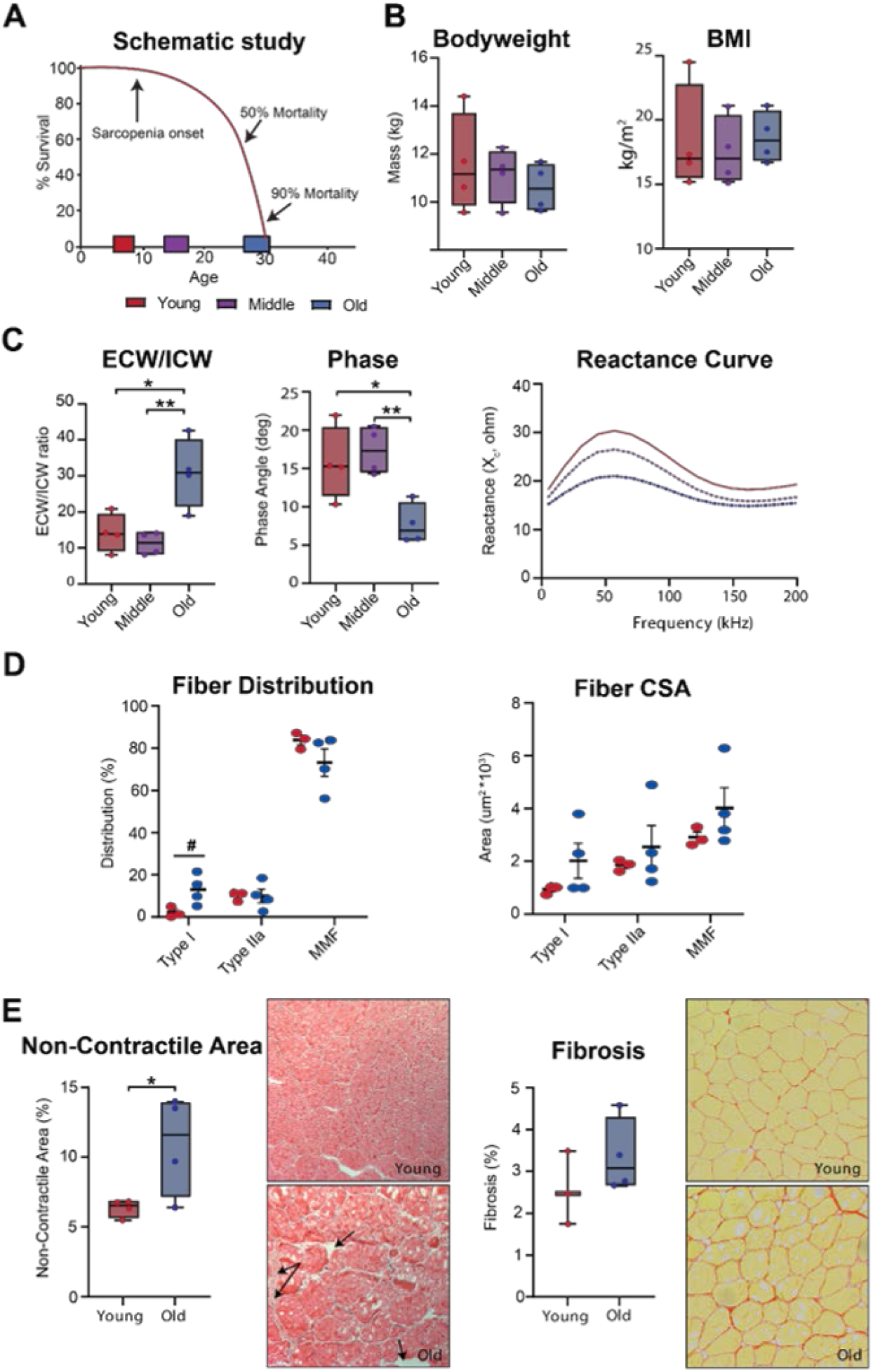
Sarcopenia is associated with changes in muscle composition. (A) Schematic showing the age of rhesus monkeys used in this study relative to a generalized survival curve. (B) Body weight and body mass index (BMI) for young (red) middle-age (purple), and old (blue) monkeys (n=4 per group). (C) Electrical properties of the upper leg (determined by S-BIS), extracellular to intracellular water ratios, phase angle, and reactance curves. (D) Distribution of fiber types vastus laterals biopsies detected using antibodies specific to type I or type II myosin heavy chain. (E) Quantification of non-contractile content (black arrows) in hematoxylin and eosin stained sections (left) and fibrotic content (right) as detected by picrosirius expressed as percent area of tissue section (4-5 images per animal). Data shown as median +/- IQR, #p<0.1, * p>0.05, ** p<0.01.

### Proteomic Analysis

To access age-related changes in the skeletal muscle proteome, proteins were extracted from the vastus lateralis from young, middle-age, and old rhesus monkeys and subject to a bottom-up proteomics analysis. Given the homology between genomes and redundancy of annotations of the UniProt *Macaca mulatta* genome, proteins were identified by searching mass spectrometry data against the UniProtKB/Swiss-Prot *Homosapien* genome.^50^ In this study, 2377 proteins were identified (Table S1) and significantly altered proteins were determined for each age group, revealing a symmetrical distribution of altered proteins across all comparisons (**Figure 3A**). Comparing old to young animals, 198 proteins were found to be significantly altered. Comparing middle-age to young animals, 647 proteins were found to be significantly altered. Comparing old to middle-age animals, 148 proteins were found to be significantly altered. There was a striking non-linearity observed in the changes of many proteins related to muscle structure as a function of age, resulting in profiles from old and young animals being more similar compared to middle-age animals to each other than either were to the profiles from middle-aged animals (**Figure 3B-E**). These including the actin, myosin, integrin, and collagen proteins. This observation suggests that the majority of muscle mass is lost between 15-28 years in our NHP model, and matches longitudinal studies indicating that vulnerability to sarcopenia begins at ∼16 years of age.^19^

**Figure 3.**
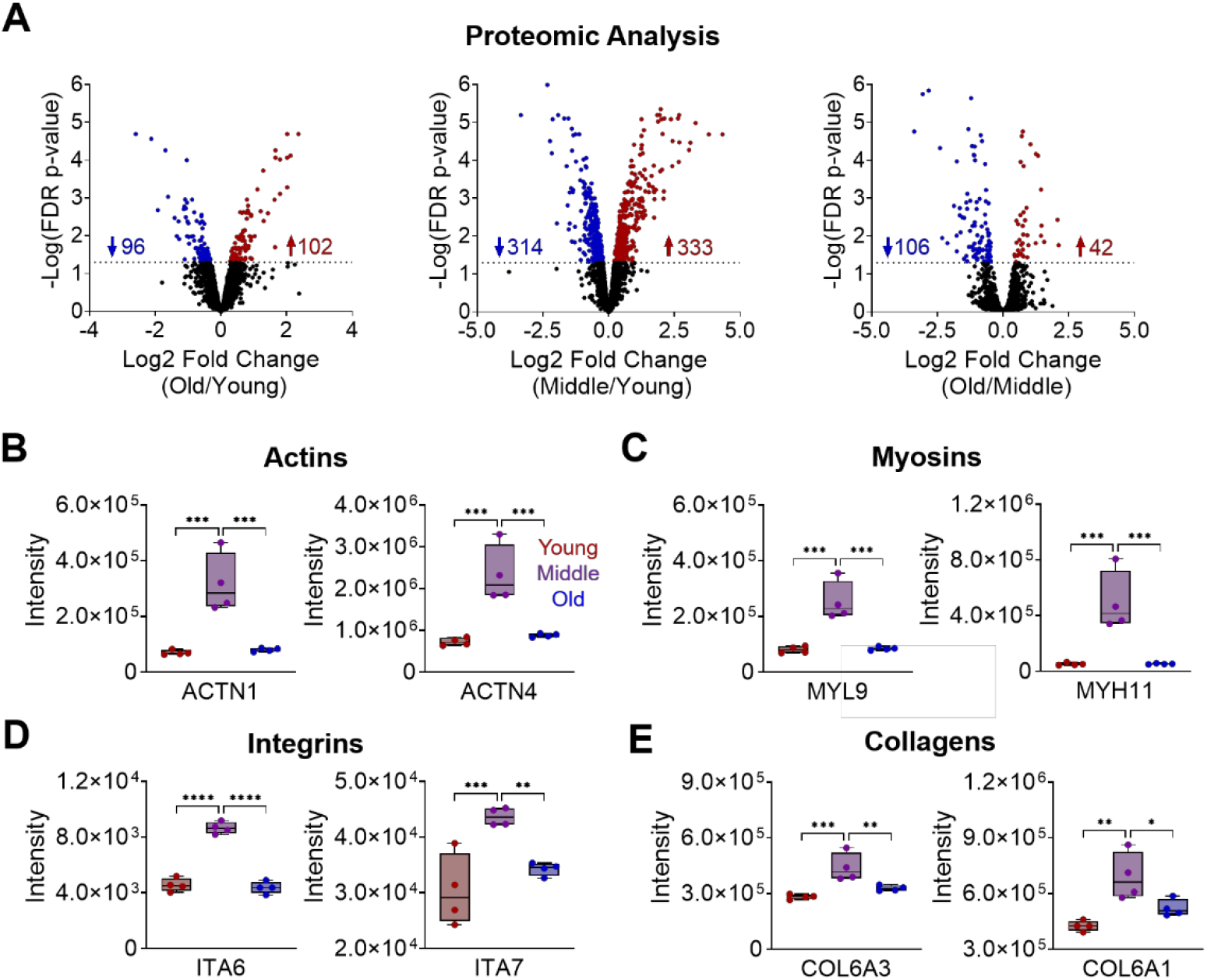
Proteomic analysis of aging rhesus monkey skeletal muscle. Proteins where extracted from the vastus lateralis from young, middle-age, and old rhesus monkeys (n=4 per group), enzymatically digested, and subjected to mass spectrometry analysis (Table S1). Volcano plots showing significantly (adjusted p-value < 0.05) altered proteins for ((A) old relative to young (left,) middle-age relative to young (middle), and old relative to middle-age animals (right). Relative levels of muscle structure proteins including (B) actins, (C) myosins, (D) integrins, and (E) collagens. *p<0.01, **p<0.001, ***p<0.0001, and ****p<0.0001. ACTN1=alpha-actinin-1, ACTN4=alpha-actinin-4, MYL9=myosin regulatory light polypeptide 9, MYH11=myosin-11, ITA6=integrin alpha-6, ITA7= integrin alpha-6, COL6A3=collagen alpha-3 (VI) chain, and COL6A1=collagen alpha-1 (VI) chain.

To gain insight into biological functions impacted by age, altered proteins were subject to GO term analysis (**Figure 4A**, Table S2). Among the top ranked enriched biological functions were adenosine triphosphate (ATP) metabolic process (GO: 0046034), adenosine monophosphate (AMP) metabolic process (GO: 0046033), and fatty acid beta oxidation (GO: 0006635). Shown are selected proteins mapped to ATP metabolic process (GO: 0046034) including ATP synthase subunit d, mitochondrial (ATPD), succinate dehydrogenase [ubiquinone] cytochrome b small subunit, mitochondrial (SDHD), and cytochrome b-c1 complex subunit 8 (QCR8), all of which show higher expression in old animals (**Figure 4B**). AMP metabolic process (GO: 0046033) was also a prevalent altered biological process, driven by lower AMP deaminase 1 (AMPD), adenylosuccinate (PUR8) and adenylosuccinate synthetase isozyme 1 (PURA1) in both middle-age and old animals (**Figure 4C**). Fatty acid beta oxidation (GO:0006635) was also found to be altered in middle-age and old animals and include higher enoyl-CoA delta isomerase 1, mitochondrial (ECI1), an enzyme that changes the position of double bonds in fatty acids joined to CoA, and acetyl-coenzyme. A carboxylase carboxyl transferase subunit alpha 2 (THIM) that catalyzes the last step in beta oxidation, and lower ethylmalonyl-CoA decarboxylase (ECHD1) an ethylmalonyl-CoA decarboxylase (**Figure 4D**).

**Figure 4.**
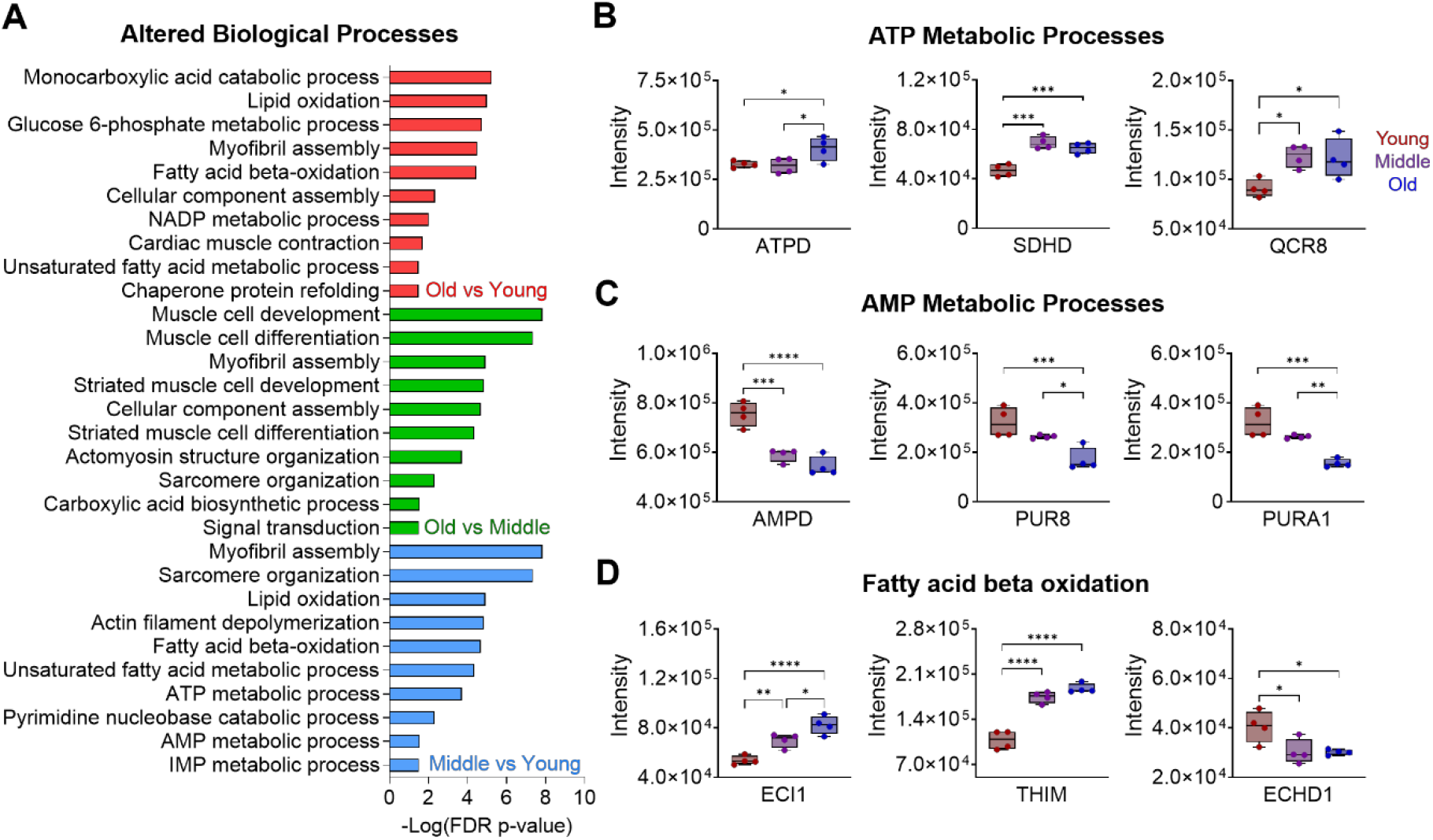
Gene Ontology (GO) term enrichment analysis of altered proteins from aging rhesus monkey skeletal muscle. Significantly altered proteins (Table S1) were subject to a GO term analysis to identify associated biological processes (Table S2). Shown are relative peak intensities for selected proteins related to selected biological functions for young, middle, and old rhesus monkeys (n=4 per group): (C) ATP metabolic process (GO: 0046034), (D) AMP metabolic process (GO: 0046033), and (E) fatty acid beta oxidation (GO:0006635). *p<0.01, ** p<0.001, ***p<0.0001, and ****p<0.0001. ITA6=integrin alpha-6, ITA7= integrin alpha-6, COL6A1=collagen alpha-1 (VI) chain, COL6A3=collagen alpha-3 (VI) chain, MYL9=myosin regulatory light polypeptide 9, MYH11=myosin-11, ACTN1=alpha-actinin-1, ACTN4=, ATPD = ATP synthase subunit d, mitochondrial, DHSD=succinate dehydrogenase [ubiquinone] cytochrome b small subunit, mitochondrial, QCR8 = cytochrome b-c1 complex subunit 8 (QCR8), AMPD= AMP deaminase 1, PUR8 = adenylosuccinate, PURA1 = adenylosuccinate synthetase isozyme 1, ECI1 = enoyl-CoA delta isomerase 1, mitochondrial, THIM = acetyl-coenzyme A carboxylase carboxyl transferase subunit alpha 2, and ECHD1=ethylmalonyl-CoA decarboxylase.

Skeletal muscle contractility is highly dependent on the availability of ATP. Previous studies have attributed lower levels of ATP in aging skeletal muscle to dysfunctional mitochondria.^51–53^ Interestingly, Pugh and co-authors describe that this shift in energy metabolism precedes sarcopenia pathophysiology, highlighting the feasibility and critical need to identify molecular targets that can be used to attenuate and/or monitor disease progression.^16^ In our analysis of the altered proteome, we observed higher expression of multiple enzymes involved in the production of ATP indicating an age-related deficiency in the present NHP model. Additionally, we observed multiple downregulated enzymes related to the production of AMP, suggesting that efforts in old animals are primarily directed to the production and conservation of ATP as opposed to production of AMP or purine nucleotide precursors. Further, we observed alterations in enzymes involved in fatty acid beta oxidation, a critical mitochondrial pathway involved in the production of ATP from lipid fuel sources. Taken together with changes in lipid processing enzymes, these data support previous findings in the literature that describe age-related ATP deficiency, and identify protein biomarkers related to altered fatty acid beta oxidation in sarcopenia.

### Metabolomic Analysis

Metabolites represent the end point of the ‘omics cascade and provide a complimentary approach to the proteomic analysis in defining the molecular landscape of muscle aging. Relative metabolite levels were evaluated in the same specimens of vastus lateralis from young, middle-age, and old rhesus monkeys. Mass spectrometric analysis allowed for detection of 8974 metabolic features, 725 of which were identified by fragmentation matching against publicly available spectral libraries (Table S3). Significantly altered metabolites were determined between age groups, revealing a symmetrical distribution of altered metabolites across all comparisons (**Figure 5A**). Comparing old to young animals, 81 metabolites were significantly altered. Comparing middle-aged to young animals, 75 metabolites were significantly altered. Comparing old to middle-aged animals, 72 metabolites were significantly altered. Among these altered metabolites were those involved in skeletal muscle energy metabolism including carnitine, trimethyllysine, and phosphocreatine, all observed to be lower in both middle-age and old animals (**Figure 5B**). phosphatidylinositol 38:4 (PI 38:4) was lower in both middle-age and old animals (**Figure 5C**). Lastly, differences in levels of acylcarnitine involved in fatty acid beta oxidation were also altered (**Figure 5D**); palmitoylcarnitine and arachidonoylcarnitine were higher in old animals relative to young animals. Integrating our proteomic and metabolomic changes together, we observed a concerted age-related dysregulation of fatty acid beta oxidation (**Figure 5E**).

**Figure 5.**
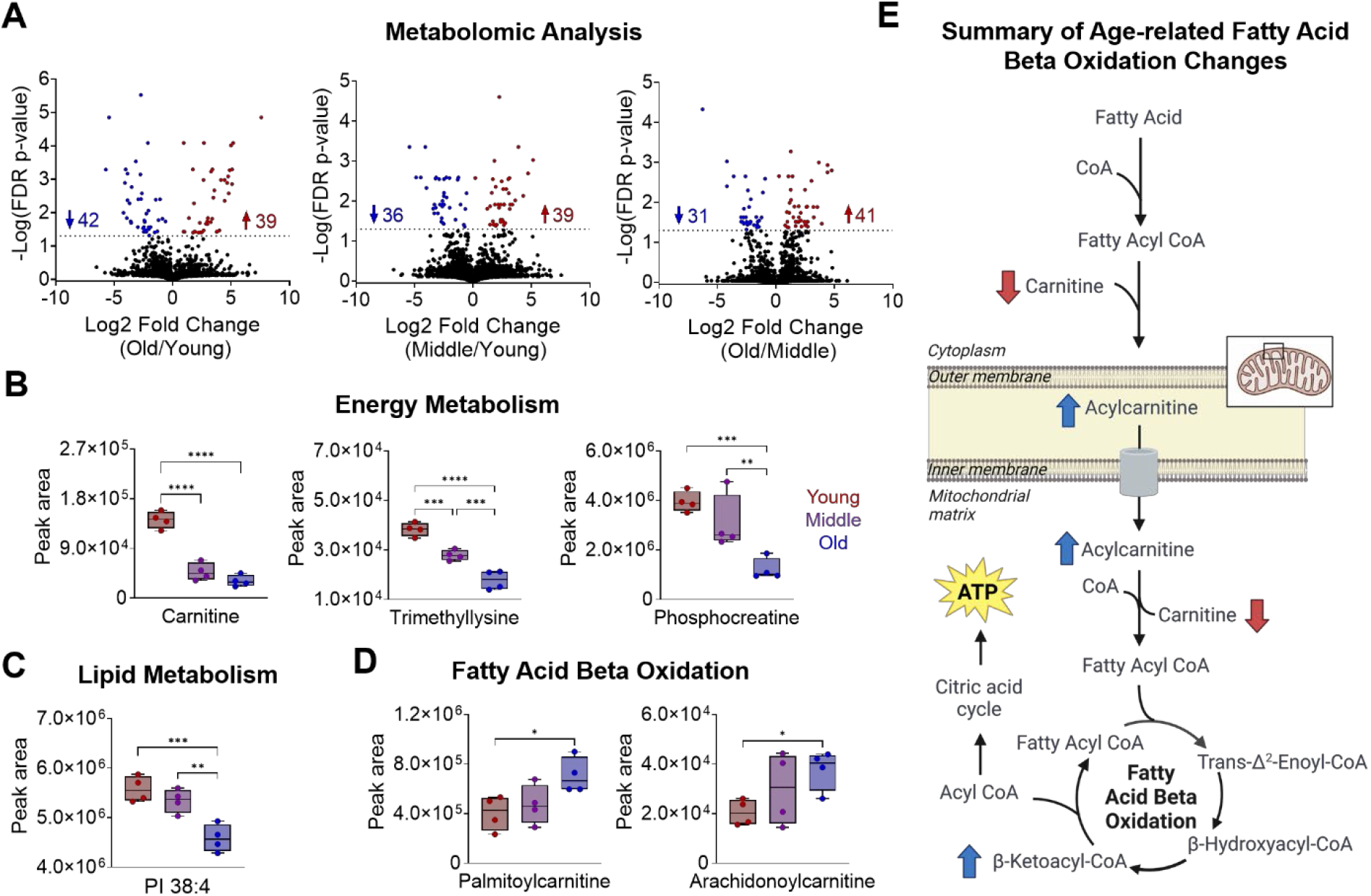
Metabolomic analysis of aging rhesus monkey skeletal muscle. Metabolites were extracted from the vastus lateralis from young, middle-age, and old rhesus monkeys (n=4 per group) and subject to mass spectrometry analysis (Table S3). Volcano plots showing significantly (adjusted p-value < 0.05) altered metabolites for (A) old relative to young (left,) middle-age relative to young (middle), and old relative to middle-age animals (right). Shown are the relative peak areas for selected metabolites related to (B) energy metabolism (carnitine (left), trimethyllysine (middle), and phosphocarnitine (right)), (C) lipid metabolism (phosphatidylinositol 38:4 (PI 38:4)), (D) fatty acid beta oxidation (palmitoylcarnitine (left) and arachidonoylcarnitine (right). (E) Summary of fatty acid beta oxidation changes observed with age. *p<0.01, **p<0.001, ***p<0.0001, and ****p<0.0001.

In the present study, we observed age-related alterations in metabolites largely associated with energy metabolism. Carnitine is essential in the translocation of fatty acids into the mitochondria for fatty acid beta oxidation. carnitine is largely synthesized in the liver, kidneys, and brain, transported in the serum, and stored in skeletal muscle until needed.^54^ Previous reports have noted low serum levels of carnitine in sarcopenia, suggesting that its production or transport efficiency may decrease with age or that it is being sequestered.^55^ Interestingly, carnitine can be synthesized by the trimethylation of lysine ^_^ an important post-translationally modified amino acid with an important function not only in carnitine biosynthesis, but also regulation of histones and downstream epigenetic regulation. Given that trimethyllysine was also observed to be lower, the results of this study suggest that there may also be a dysregulation of lysine methylation, thereby contributing to lower levels of carnitine. Moreover, phosphocreatine is stored in skeletal muscle under normal conditions to provide an accessible supply of phosphate for the production of ATP.^11^ Apart from carnitine metabolism, lower levels of phosphocreatine were observed in old animals. With this, it is reasonable to infer that lack of phosphocreatine would further blunt the ability to efficiently produce enough ATP to maintain muscle strength and contractile function. The higher palmioylcarnitine and arachidonoylcarnitine in old animals could reflect a failure to efficiently use lipid fuel.^56^ Higher levels of tissue resident and circulating acylcarnitines have being linked to defects in skeletal muscle beta oxidation and have been observed in muscle with sarcobesity and lipid stress.^57, 58^ Lastly, phosphatidylinositols are important lipids known to be incorporated into the membranes of several organelles, including the mitochondria.^59^ Overall, the results of our metabolomic analysis support previous findings of age-related changes in energy metabolism in skeletal muscle and also provide concerted protein-metabolite perturbations suggestive of dysregulation of fatty acid beta oxidation in sarcopenia.

### Integration of Proteomic and Metabolomic Analyses

An unbiased, metabolomics and proteomics pathway analysis was performed to identify altered pathways in our NHP model of sarcopenia (**Figure 6**, Table S4). Comparing old to young animals, fatty acid degradation (Figure S1) and fatty acid elongation pathways (Figure S2) were upregulated whereas purine metabolism (Figure S3) and fatty acid biosynthesis pathways (Figure S4) were downregulated (**Figure 6A**). Comparing middle-age to young animals, fatty acid degradation (Figure S1), fatty acid elongation (Figure S2), focal adhesion (Figure S5), and regulation of the actin cytoskeleton (Figure S6) were upregulated whereas glycolysis (Figure S7), purine metabolism (Figure S3), and fatty acid biosynthesis (Figure S4) were downregulated (**Figure 6B**). Comparing old to middle-age animals, starch and sucrose metabolism (Figure S8) were upregulated whereas fatty acid biosynthesis (Figure S4), muscle contraction (Figure S9), regulation of actin cytoskeleton (Figure S5), and focal adhesion (Figure S6) were downregulated (**Figure 6C**). Together, these data describe age-responsive pathways that occur in sarcopenia and broadly include changes in muscle structure and contraction and alterations in lipid, carbohydrate, and purine metabolism (**Figure 6D**).

**Figure 6.**
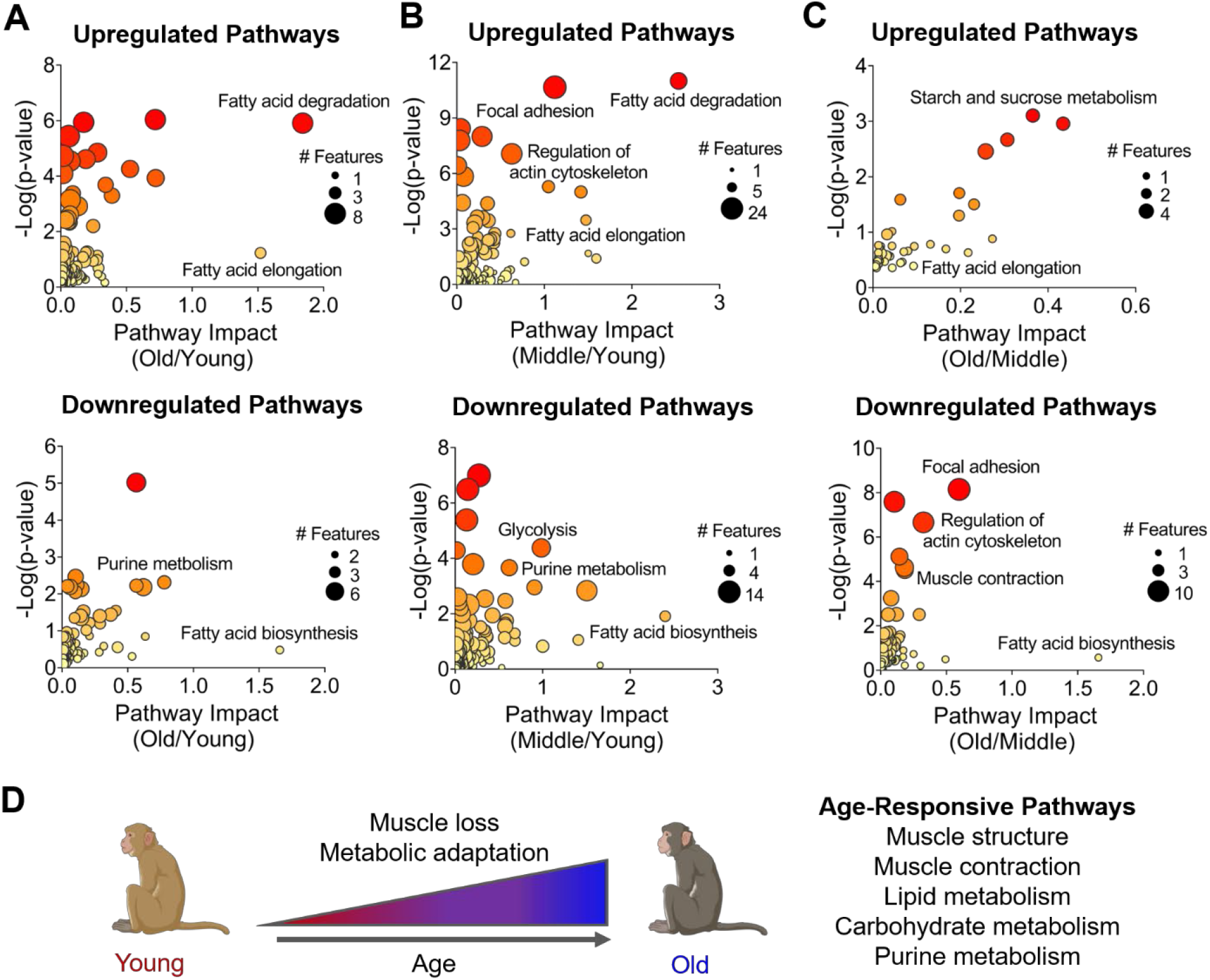
Integrated protein and metabolite pathway analysis. Altered proteins and metabolites were subject to KEGG pathway analysis (Table S4). Shown are bubble plots depicting top-ranked up (top) and downregulated (bottom) pathways for (A) old relative to young, (B) middle-age relative to young and (C) old relative to middle-age animals. Bubble size is directly related to the number of significantly altered protein and metabolite (features). Color gradient is related to significance of pathway enrichment (low (yellow) to high (red)). (D) Schematic summarizing the most prominent age-responsive pathway changes.

The results of this study provide a foundation for mechanistic insight into the molecular changes that occur in aging skeletal muscle. Nearly all proteins mapped to the focal adhesion and regulation of the actin cytoskeleton pathways were lower in old compared to middle-age animals. Notably, many of these proteins are involved in the support and organization of skeletal muscle.^60–62^ A dysregulation in cytoskeleton dynamics in old animals was also indicated by alteration of integrins, laminins and serine/threonine-protein kinases.^60, 63, 64^ Many pathways broadly associated with changes in energy metabolism were found to be altered with age; a shift that is consistent with numerous reports in the literature. The decrease in purine metabolism would presumably have a downstream effect on nucleotide synthesis, and could impact production of metabolic cofactors, such as NAD and NADP, in addition to energy storing nucleotide triphosphates that are involved in fuel mobilization and intracellular signaling.^65^ Changes in lipid degradation and synthetic pathways point to dysfunction in lipid handling with age, potentially contributing to energetic deficit. Lipids also play a role in critical cellular functions such as energy stores, structural membranes, secretory vesicles, and as second messengers involved in nutrient and inflammatory responses.^66–68^ Collectively, these data provide additional knowledge to aid in the better understanding of the molecular determinants underlying sarcopenia.

## Conclusions

This study explores altered proteins and metabolites in the vastus lateralis of aging rhesus monkeys. Mass spectrometry analysis of the proteome and metabolome identified concerted sarcopenia-associated perturbations. In our proteomic analysis, expression of proteins related to adenosine triphosphate and adenosine monophosphate metabolism as well as fatty acid beta oxidation were all age-responsive. In our metabolomic analysis, metabolites involved in energy metabolism and fatty acid beta oxidation were also age-sensitive. Pathway analysis of aggregate proteome and metabolome data altered between age groups confirmed a dysregulation of energy metabolism and further identified muscle structure and composition as well as lipid, carbohydrate, and purine metabolism increasing age-responsive process. Collectively, this study identified new metabolic signatures in sarcopenia to uncover underlying molecular mechanism in sarcopenia.

## Supporting information

Table S1 Proteomics

Table S2 Altered Proteins GO Term Analysis

Table S3 Metabolomics

Table S4 Pathway Analysis

## Data Availability

All data contained within this manuscript is available upon request of the corresponding authors. All mass spectrometry data have been deposited to the MASSive data repository with the dataset identifier MSV000091537.

## Acknowledgements

This research is supported by NIH R01 HL109810-06 (Y.G.). Y.G. acknowledges NIH R01 GM125085, R01 HL096971, and S10 OD018475. D.S.R. acknowledges the support from the American Heart Association Predoctoral Fellowship Grant (No. 832615). K.J.R acknowledges the National Science Foundation Graduate Research Fellowship Program (No. DGE-1747503). RMA and RJC acknowledge NIH R01 AG040178 and R01 AG074503. The Wisconsin National Primate Research Center is supported by P51OD011106 and Research Facilities Improvement Programs RR15459-01 and RR020141-01. The study was conducted with the use of resources and facilities at the Geriatric Research Education and Clinical Center in the William S. Middleton Memorial Veterans Hospital, Madison WI.

## Conflict of Interest

The authors declare no conflicts of interest.

## Notes

ftp://massive.ucsd.edu/MSV000091537/

### Competing Interest Statement

The authors have declared no competing interest.

