## Supporting Information for "Mass spectrometry-based multi-omics identifies metabolic signatures of sarcopenia in rhesus monkey skeletal muscle"

#### Table of Contents


| | | |
|---|---|---|
| Table S1 | Proteomics | |
| Table S2 | GO Term Analysis | |
| Table S3 | Metabolomics | |
| Table S4 | Pathway Analysis | |
| Figure S1 | Features mapped to fatty acid degradation | S-3 |
| Figure S2 | Features mapped to fatty acid elongation | S-4 |
| Figure S3 | Features mapped to purine metabolism | S-5 |
| Figure S4 | Feature mapped to fatty acid biosynthesis | S-6 |
| Figure S5 | Features mapped to focal adhesion | S-7 |
| Figure S6 | Features mapped to regulation of actin cytoskeleton | S-8 |
| Figure S7 | Features mapped to glycolysis | S-9 |
| Figure S8 | Features mapped to starch and sucrose metabolism | S-10 |
| Figure S9 | Features mapped to muscle contractility | S-11 |

### Fatty acid degradation

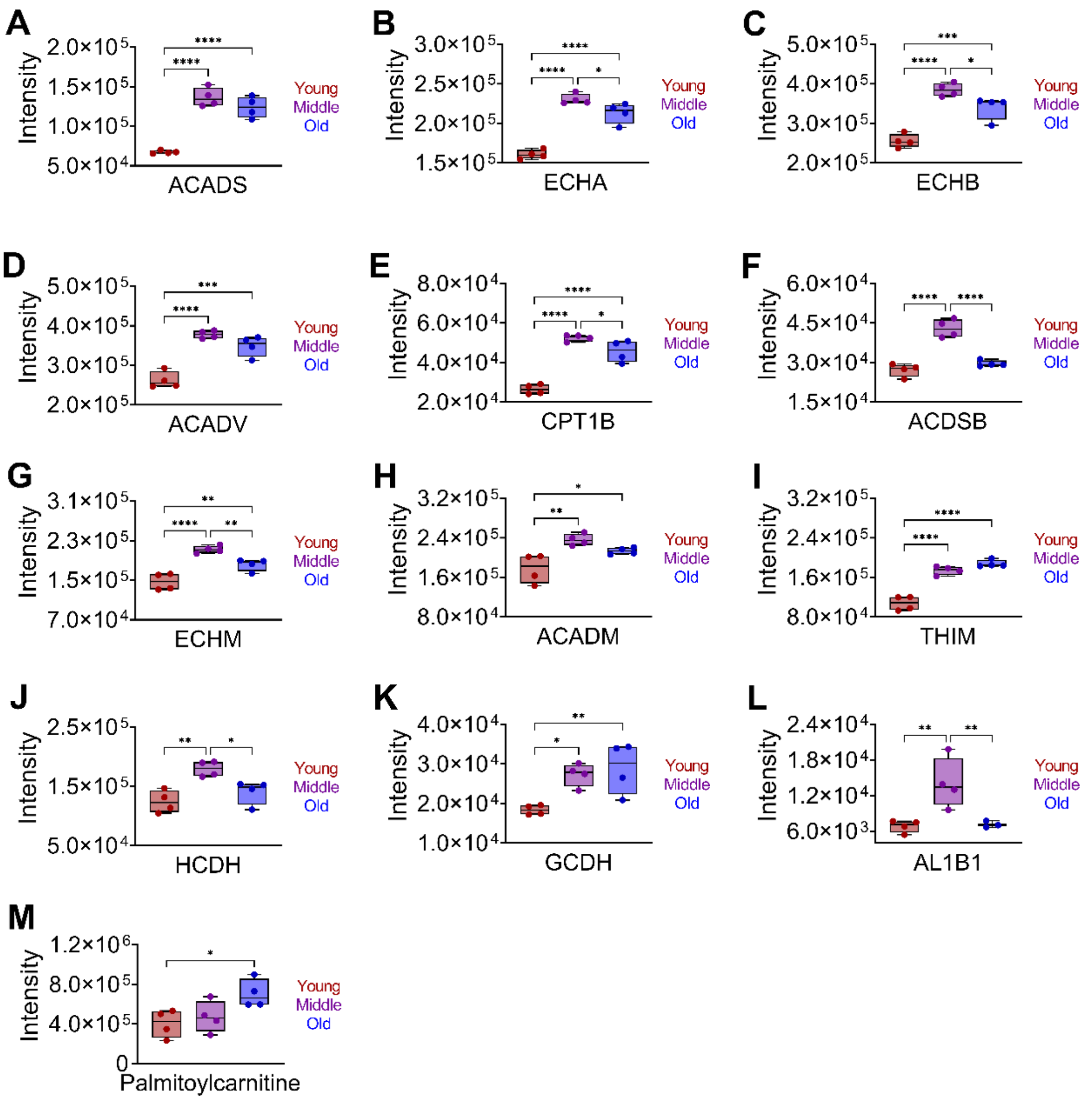


**Figure S1. Features mapped to fatty acid degradation.** Pathway analysis for significantly altered (adjusted p-value $\leq$ 0.05) features (Table S4) for young, middle-age, and old (n=4 per group) rhesus monkeys. Shown are relative peak intensities for features mapped to fatty acid degradation. (A) ACADS=short-chain specific acyl-CoA dehydrogenase, mitochondrial, (B) ECHA = **trifunctional enzyme subunit alpha, mitochondrial,** (C) **ECHB = trifunctional enzyme subunit beta, mitochondrial, (D) ACADV = very long-chain specific acyl-CoA dehydrogenase, mitochondrial,** (E) **CPT1B =** Carnitine O-palmitoyltransferase 1, muscle isoform, (F) ACDSB = **short/branched chain specific acyl-CoA dehydrogenase, mitochondrial,** mitochondrial, (G) ECHM = enoyl-CoA hydratase, mitochondrial, (H) ACADM = medium-chain specific acyl-CoA dehydrogenase, mitochondrial, (I) THIM = 3-ketoacyl-CoA thiolase, mitochondrial, (J) **HCDH =** hydroxyacyl-coenzyme A dehydrogenase, (K) GCDH = glutaryl-CoA dehydrogenase, mitochondrial, AL1B1= aldehyde dehydrogenase X, mitochondrial, (L) ECI1 = **Enoyl-CoA delta isomerase 1, mitochondrial, and** (M) **palmitoylcarnitine.** *****p<0.01, **p$\leq$0.001, ***p$\leq$0.0001, and ****p$\leq$0.0001.

### Fatty acid elongation

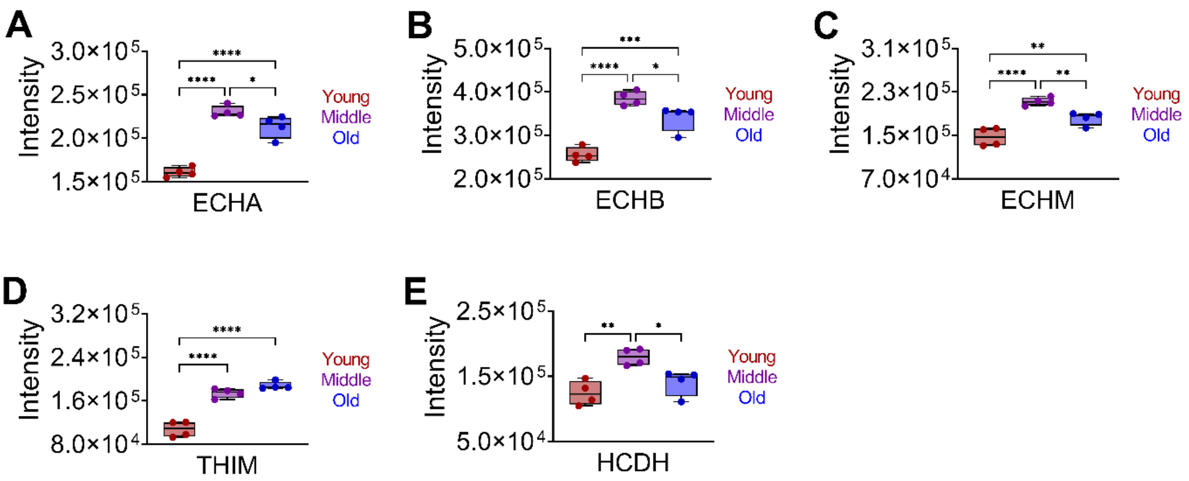


**Figure S2. Features mapped to fatty acid elongation.** Pathway analysis for significantly altered (adjusted p-value $\le$ 0.05) features (Table S4) for young, middle-age, and old (n=4 per group) rhesus monkeys. Shown are relative peak intensities for features mapped to fatty acid elongation. (A) ECHA = **trifunctional enzyme subunit alpha, mitochondrial,** (B) **ECHB = trifunctional enzyme subunit beta, mitochondrial,** (A) ACADS=short-chain specific acyl-CoA dehydrogenase, mitochondrial, (C) ECHM = enoyl-CoA hydratase, mitochondrial**,** (D) THIM = 3-ketoacyl-CoA thiolase, mitochondrial, and (E) **HCDH =** hydroxyacyl-coenzyme A dehydrogenase. *****p<0.01, **p$\le$0.001, ***p$\le$0.0001, and ****p$\le$0.0001.

### Purine metabolism

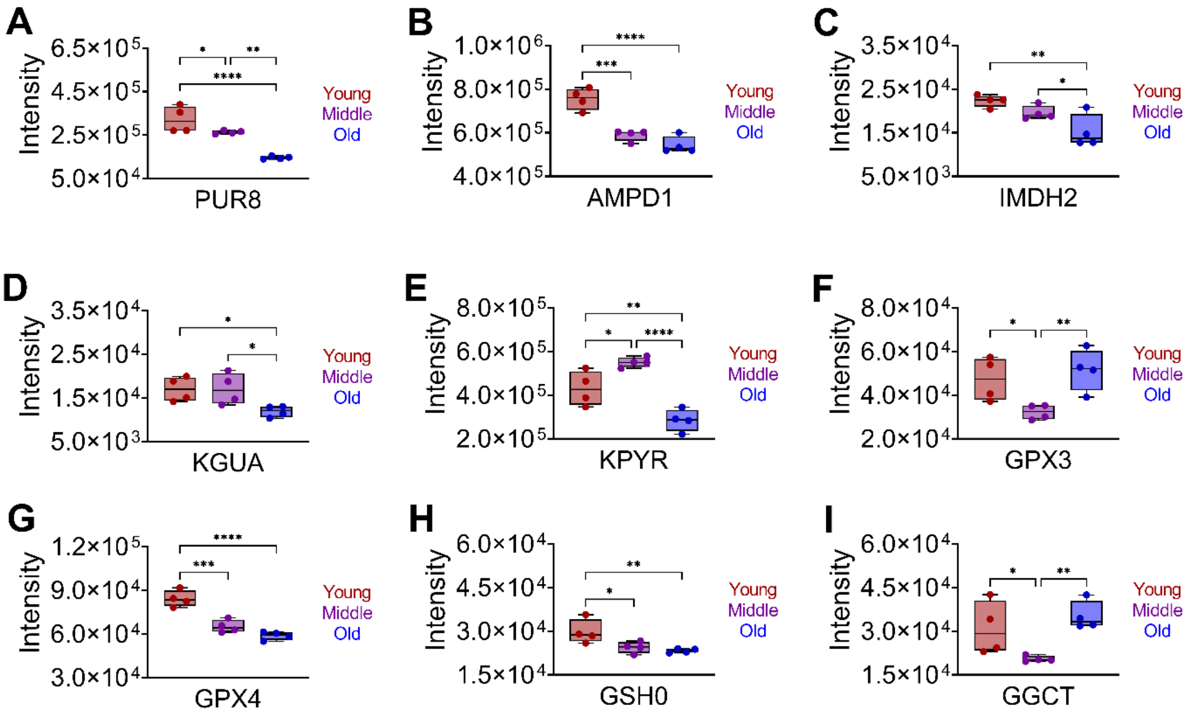


**Figure S3. Features mapped to purine metabolism.** Pathway analysis for significantly altered (adjusted p-value $\leq$ 0.05) features (Table S4) for young, middle-age, and old (n=4 per group) rhesus monkeys. Shown are relative peak intensities for features mapped to purine metabolism. (A) PUR8 = **adenylosuccinate lyase, (B) AMPD1 = AMP deaminase 1, (C) IMDH2 = inosine-5'-monophosphate dehydrogenase 2, (D) KGUA = guanylate kinase, (E) PYR = pyruvate kinase PKLR, (F) GPX3 =** glutathione peroxidase 3, (G) GPX4 = phospholipid hydroperoxide glutathione peroxidase, (H) GSH0 = glutamate--cysteine ligase regulatory subunit, and (I) GGCT = gamma-glutamylcyclotransferase. *****p<0.01, **p$\leq$0.001, ***p$\leq$0.0001, and ****p$\leq$0.0001.

### Fatty acid biosynthesis

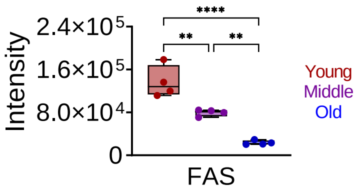


**Figure S4. Feature mapped to fatty acid biosynthesis.** Pathway analysis for significantly altered (adjusted p-value $\leq$ 0.05) features (Table S4) for young, middle-age, and old (n=4 per group) rhesus monkeys. Shown are relative peak intensities for features mapped to fatty acid biosynthesis. FAS = fatty acid synthase. **p$\leq$0.001 and ****p$\leq$0.0001.

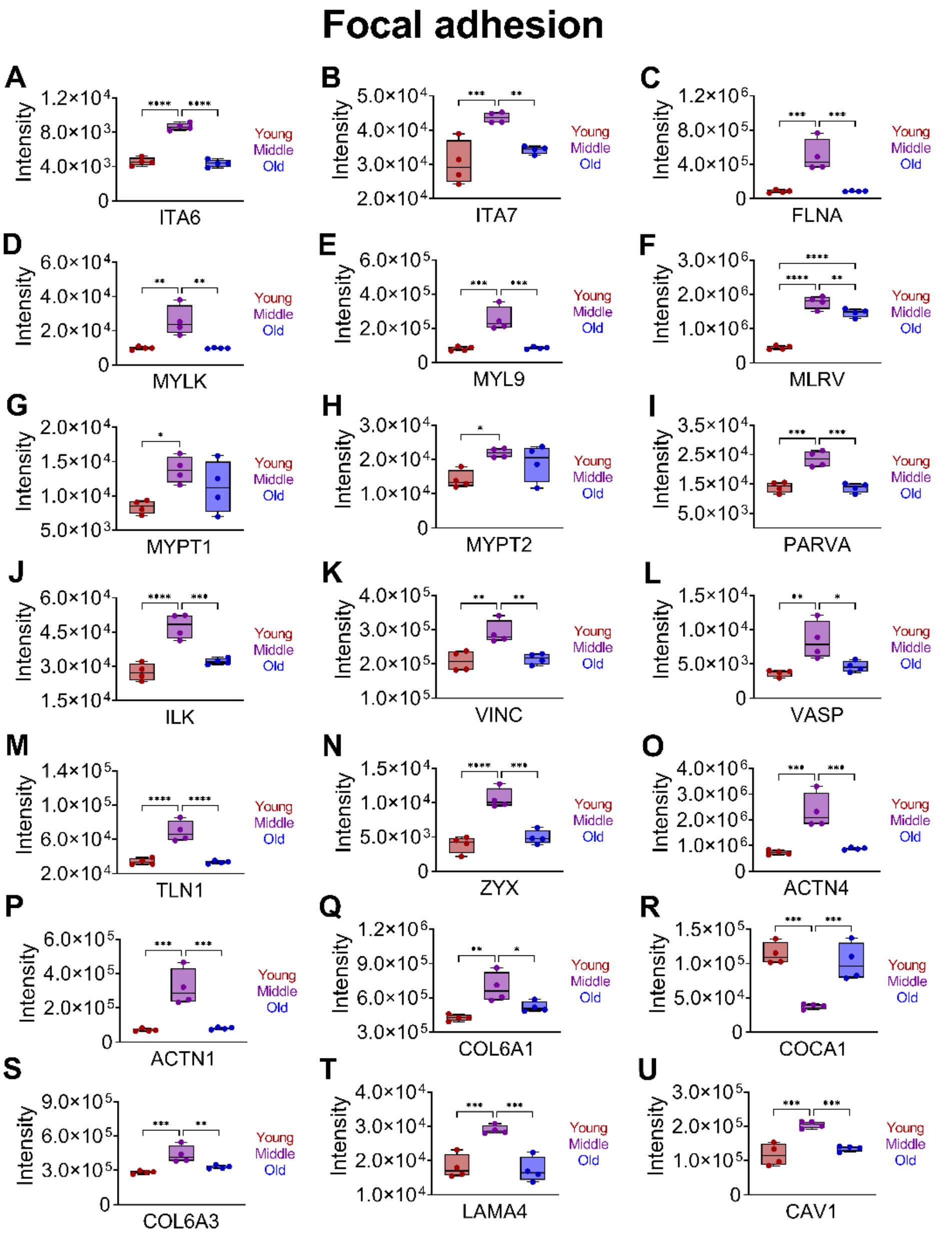


**Figure S5. Features mapped to focal adhesion.** Pathway analysis for significantly altered (adjusted p-value $\leq$ 0.05) features (Table S4) for young, middle-age, and old (n=4 per group) rhesus monkeys. Shown are relative peak intensities for features mapped to focal adhesion. (A) ITA6 = integrin alpha-6, (B) ITA7 = integrin alpha-7, (C) filamin-A, (D) MYLK= myosin light chain kinase, smooth muscle, (E) MYL9 = myosin regulatory light polypeptide 9, (F) MLRV = myosin regulatory light chain 2, ventricular/cardiac muscle isoform, (G) MYPT1 = protein phosphatase 1 regulatory subunit 12A, (H) MYPT2 = protein phosphatase 1 regulatory subunit 12B, (I) PARVA = alpha-parvin, (J) ILK = integrin-linked protein kinase, (K) VINC = vinculin, (L) VASP = vasodilator-stimulated phosphoprotein, (M) TLN1 = talin-1, (N) ZYX = zyxin, (O) ACTN4 = alpha-actinin-4, (P) ACTN1 = alpha-actinin-1, (Q) CO6A1 = collagen alpha-1(VI) chain, (R) COCA1 = collagen alpha-1(XII) chain, (S) COL6A3 = collagen alpha-3(VI) chain, (T) LAMA4 = laminin subunit alpha-4, and (U) CAV1 = caveolin-1.* $p<0.01$, **$p\leq0.001$, and ***$p\leq0.0001$.

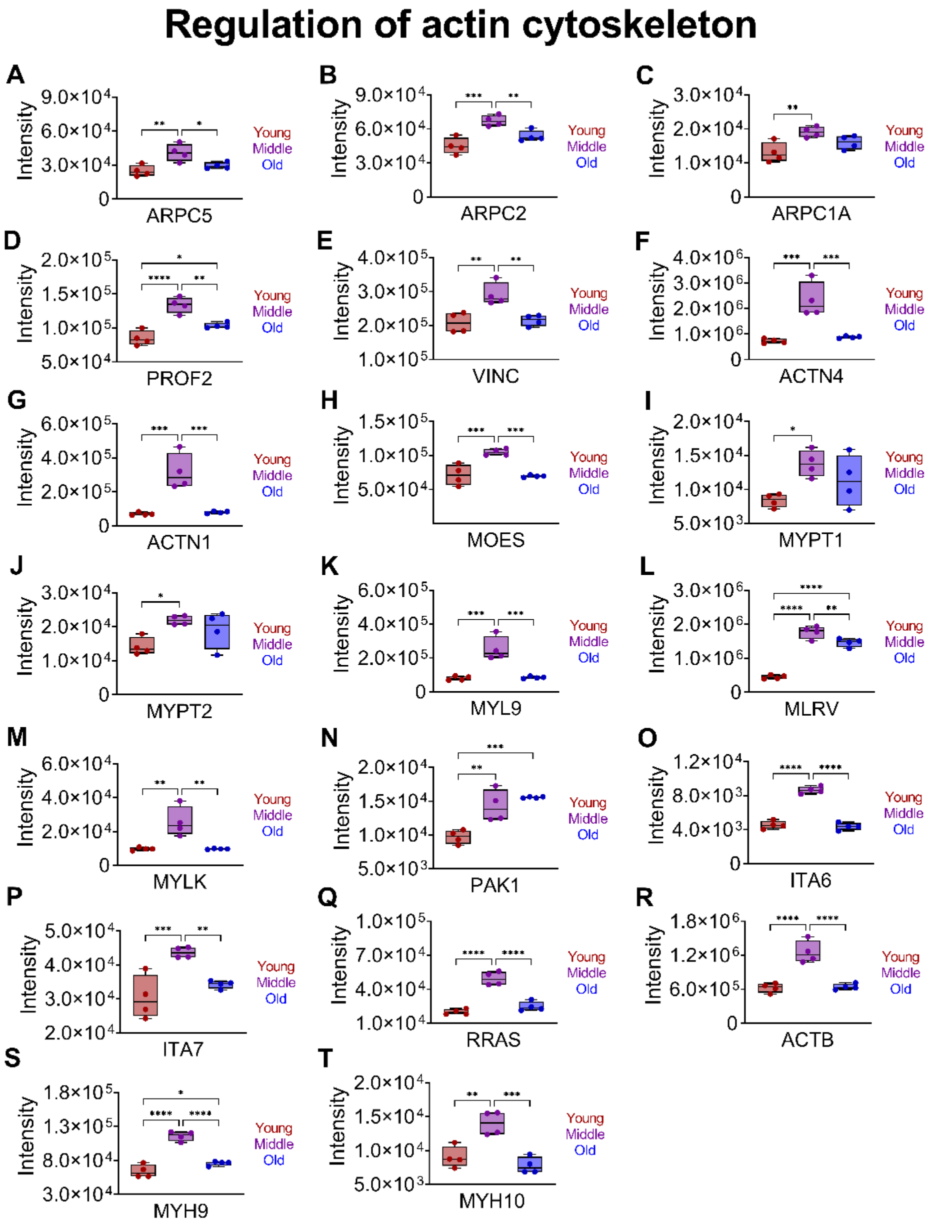


**Figure S6. Features mapped to regulation of actin cytoskeleton.** Pathway analysis for significantly altered (adjusted p-value $\leq$ 0.05) features (Table S4) for young, middle-age, and old (n=4 per group) rhesus monkeys. Shown are relative peak intensities for features mapped to regulation of actin cytoskeleton. (A) ARPC5 = actin-related protein 2/3 complex subunit 5, (B) ARPC2 = actin-related protein 2/3 complex subunit 2, (C) ARPC1A = actin-related protein 2/3 complex subunit 1A, (D) PROF2 = profilin-2, (E) VINC = vinculin, (F) ACTN4 = alpha-actinin-4, (G) ACTN1 = alpha-actinin-1, (H) MOES = moesin, (I) MYPT1 = protein phosphatase 1 regulatory subunit 12A, (J) MYPT2 = protein phosphatase 1 regulatory subunit 12B, (K) MYL9 = myosin regulatory light polypeptide 9, (L) MLRV = myosin regulatory light chain 2, ventricular/cardiac muscle isoform, (M) MYLK= myosin light chain kinase, (N) serine/threonine-protein kinase PAK1, (O) ITA6 = integrin alpha-6, (P) ITA7 = integrin alpha-7, (Q) RRAS = ras-related* protein R-Ras, (R) ACTB = actin, cytoplasmic, (S) MYH9 = myosin-9, and (T) MYH10 = myosin-10. p<0.01, **p$\leq$0.001, and ***p$\leq$0.0001.

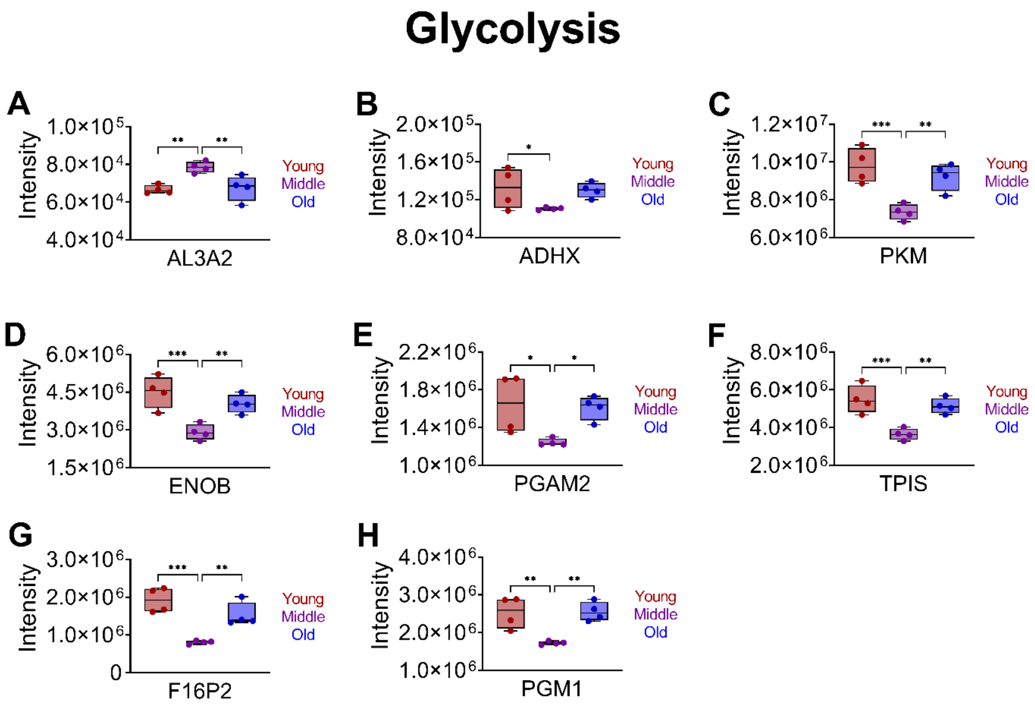


**Figure S7. Features mapped to glycolysis.** Pathway analysis for significantly altered (adjusted p-value ≤ 0.05) features (Table S4) for young, middle-age, and old (n=4 per group) rhesus monkeys. Shown are relative peak intensities for features mapped to glycolysis. (A) AL3A2 = adehyde dehydrogenase family 3-member A2, (B) ALDX = alcohol dehydrogenase class-3, (C) PKM = pyruvate kinase PKM, (D) ENOB = beta-enolase, (E) PGAM2 = phosphoglycerate mutase 2, (F) TPIS = triphosphate isomerase, (G) F16P2 = fructose-1,6-bisphosphatase isozyme 2, and (H) PGM1 = phosphoglucomutase-1. *p<0.01, **p≤0.001, and p≤0.0001***.

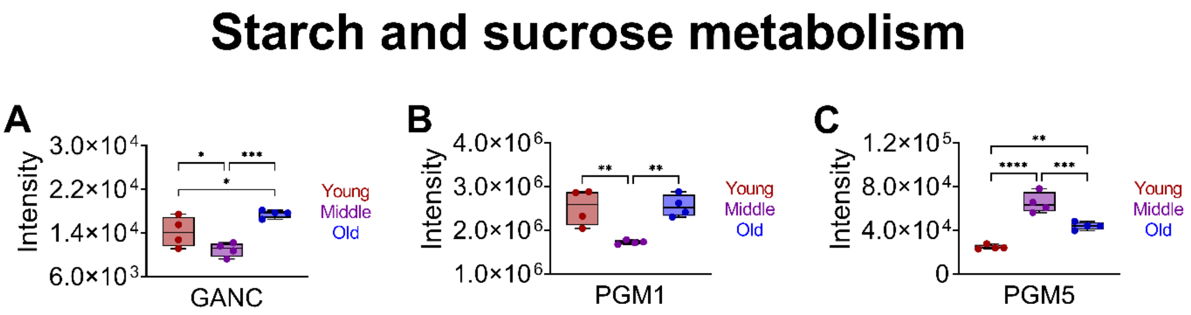


**Figure S8. Features mapped to starch and sucrose metabolism.** Pathway analysis for significantly altered (adjusted p-value $\leq$ 0.05) features (Table S4) for young, middle-age, and old (n=4 per group) rhesus monkeys. Shown are relative peak intensities for features mapped to starch and sucrose metabolism. (A) GANC = neutral alpha-glucosidase C, (B) PGM1 = phosphoglucomutase-1, and (C) PGM5 = phosphoglucomutase-like protein 5. *$p<0.01$, **$p\leq0.001$, ***$p\leq0.0001$, and ****$p\leq0.0001$.

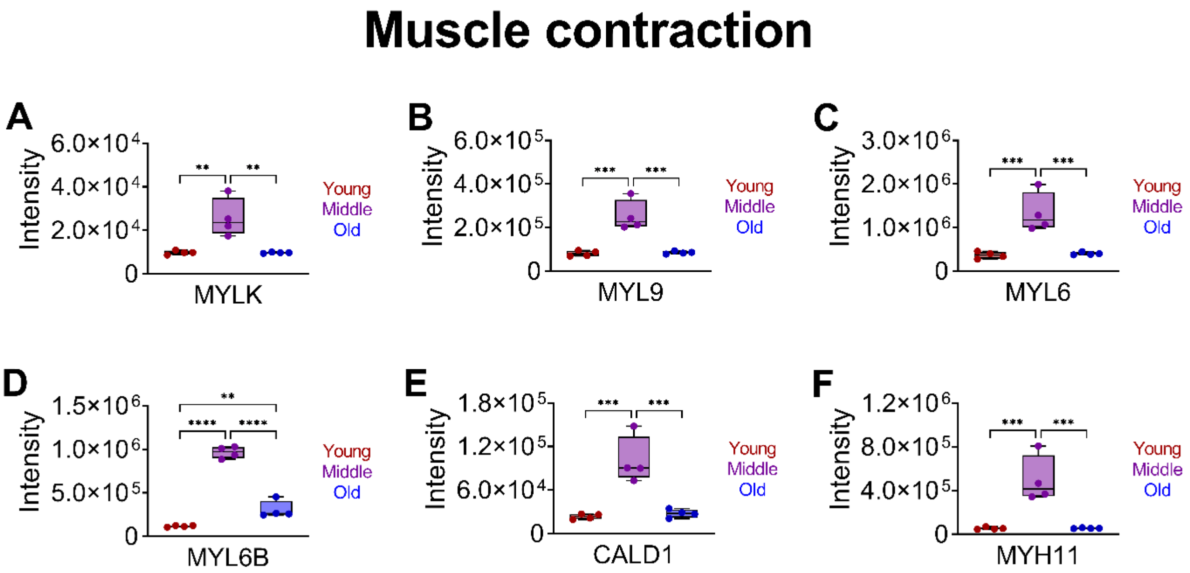


**Figure S9. Features mapped to muscle contractility.** Pathway analysis for significantly altered (adjusted p-value $\leq$ 0.05) features (Table S4) for young, middle-age, and old (n=4 per group) rhesus monkeys. Shown are relative peak intensities for features mapped to muscle contraction. (A) MYLK= myosin light chain kinase, smooth muscle, (B) MYL9 = myosin regulatory light polypeptide 9, (C) MYL6 = myosin light polypeptide 6, (D) MYL6B = myosin light chain 6B, (E) CALD1 = caldesmon, and (F) MYH-11 = myosin-11. *p<0.01, **p$\leq$0.001, ***p$\leq$0.0001, and ****p$\leq$0.0001.
